# Diverse homeostatic responses to visual deprivation by uncovering recurrent subnetworks

**DOI:** 10.1101/312926

**Authors:** Yann Sweeney, Samuel J. Barnes, Claudia Clopath

## Abstract

Multiple homeostatic plasticity mechanisms are thought to be critical for the prevention of excessively high or aberrantly low neural activity in the adult cortex. In L2/3 of adult mouse visual cortex the interplay between disinhibition and local functional interactions may support homeostatic recovery following visual deprivation. Despite blanket disinhibition only a subset of L2/3 excitatory neurons are observed to exhibit homeostatic recovery. Recovering neurons tend to be correlated with each other, forming functional networks prior to deprivation. How homeostatic recovery occurs in this way is therefore unclear, particularly in conditions of global disinhibition.

Here, we employ a computational modelling approach to investigate the origin of diverse homeostatic responses in the cortex. This model finds network size to be a critical determinant of the diverse homeostatic activity profiles observed following visual deprivation, as neurons which belong to larger networks exhibit a stronger homeostatic response. Our simulations provide mechanistic insights into the emergence of diverse homeostatic responses, and predict that neurons with a high proportion of enduring functional associations will exhibit the strongest homeostatic recovery. We test and confirm these predictions experimentally.

## INTRODUCTION

Homeostatic plasticity is widely thought to regulate neural activity. Visual deprivation paradigms are used to uncover mechanisms which mediate homeostatic plasticity in visual cortex. Numerous mechanisms have been implicated in the recovery of activity following visual deprivation, predominantly the reduction of inhibition and excitatory synaptic scaling [Rose et al., 2016, Li et al., 2014, Hengen et al., 2013, Keck et al., 2013, Kuhlman et al., 2013, Ma et al., 2013, Turrigiano et al., 1998]. Recent experiments established that homeostatic recovery from visual deprivation in layer 2/3 of adult mice is associated with a period of disinhibition, but did not observe any evidence of synaptic scaling [Barnes et al., 2015]. Surprisingly, these experiments observed that only a subset neurons recover their activity in vivo, even though all neurons undergo a blanket reduction in inhibition [Barnes et al., 2015]. It is not clear how to reconcile such global disinhibition with the observed selective homeostatic recovery within functional neural circuits. Although the impact of homeostatic plasticity on neural circuit dynamics has been widely explored in theoretical models, most require some form of synaptic scaling, and none have investigated how these diverse homeostatic responses to visual deprivation can emerge despite blanket disinhibition [Sammons et al., 2018, Barnes et al., 2017, Clopath et al., 2016, Zenke et al., 2015, Toyoizumi et al., 2014, Srinivasa and Jiang, 2013, Luz and Shamir, 2012, Vogels et al., 2011].

We use computational modelling, validated by experimental data, to suggest a mechanism underlying the diverse activity profiles of neurons following deprivation. We construct a recurrent network in which neurons form two types of networks: those driven by common visual inputs, and those driven by correlated spontaneous activity. Visual deprivation triggers inhibitory synaptic plasticity - but not synaptic scaling - within the recurrent network, consistent with observations by [Barnes et al., 2015]. Homeostatic recovery from visual deprivation in the network model is driven by the disinhibitory effect of inhibitory synaptic plasticity, which unmasks recurrent excitation in the subnetworks formed by correlated spontaneous activity. Using this network model we explore the conditions under which disinhibition successfully unmasks recurrent excitation to recover neural activity.

Crucially, for each neuron the magnitude of the homeostatic response depends on the size of the spontaneous subnetwork to which it belongs. Neurons belonging to small spontaneous subnetworks share a relatively low amount of recurrent excitation, whereas neurons belonging to large spontaneous subnetworks have stronger recurrent excitation. This increased excitation allows a much larger homeostatic response when disinhibition occurs, due to the effect of recurrent amplification previously described in computational models and in vivo [MacLean et al., 2005, Douglas et al., 1995]. Our simulation provides a mechanistic explanation for the diverse homeostatic responses to visual deprivation observed in vivo during blanket disinhibition. Our model demonstrates that disinhibition can drive homeostatic recovery in the absence of synaptic scaling, but only given sufficient recurrent excitation. Moreover, when we compare our model of homeostatic recovery to a simple model of recovery mediated by synaptic scaling, we find that scaling-mediated homeostatic responses are not subnetwork-specific, and occur uniformly across all neurons in the network. Finally, our network model predicted that neurons with a high proportion of enduring functional associations will exhibit the strongest homeostatic recovery. By imaging neurons in L2/3 of visual cortex in vivo after enucleation, we confirmed these predictions experimentally.

## MATERIALS AND METHODS

### Neuron model

We use a simple firing rate neuron model, given by the transfer function g(*x*) defined below, and as used previously by [Rajan et al., 2010, Hennequin et al., 2014].

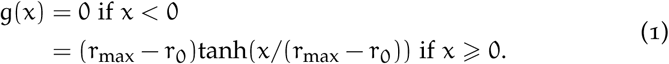

This leads to firing rates with a baseline of r_0_ and a maximum of r_max_. Following [Rajan et al., 2010], the firing rates y_i_ of neuron i driven by external input H_i_ in a network are described below.

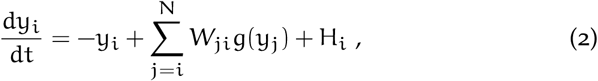

where W_ji_ is the weight of the synaptic connection from neuron j to neuron *i*.

### Recurrent network model

Our network consists of N_E_ excitatory neurons and N_I_ inhibitory neurons. The dynamics of both inhibitory (I) and excitatory (E) neurons are described by Equation 1 and Equation 2. There is dense all-to-all synaptic connectivity in the E-E, E-I and I-E populations, and no I-I connectivity. Self-connections, or autapses, are not permitted in this network. As such, W_ij_ in Equation 2 takes the form of a square matrix with size (N_E_ + N_I_)^2^, where N_E_ and N_I_ represent the number of excitatory and inhibitory neurons respectively. The strength of the inhibitory synapses are set so that inhibitory currents roughly balances excitatory currents in the network. There are 400 excitatory and 100 inhibitory neurons in the network.

### Modelling synaptic plasticity

We use the BCM learning rule to model excitatory synaptic plasticity of recurrent excitatory to excitatory (E-E) and excitatory to inhibitory (E-I) synapses [Bienenstock et al., 1982, Blais and Cooper, 2008],

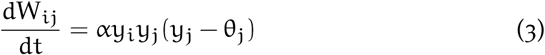

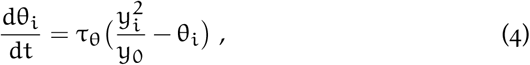

where *α* is the learning rate. θ_*i*_ refers to the sliding threshold which determines whether potentiation or depression occurs for synapses onto each neuron *i*, and which depends on the neuron’s recent postsynaptic activity, *y*_i_. τ_θ_ is the time constant at which θ_i_ is modified in order to maintain the postsynaptic firing rate at its homeostatic target, y_0_.

The BCM learning rule has both a Hebbian component and a homeostatic component. This form of plasticity is competitive and leads to the development of stimulus selectivity, as discussed in [Bienenstock et al., 1982]. In addition to the BCM learning rule, we implement synaptic pruning of weak excitatory synapses, which occurs continuously throughout development. This is implemented by deleting synapses which fall below a threshold weight set at 25% of the maximum weight.

We use a homeostatic rule to model inhibitory synaptic plasticity of recurrent inhibitory to excitatory (I-E) weights [Vogels et al., 2011],

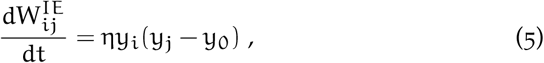
where y_0_ is the homeostatic target firing rate, η is the learning rate, and 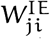 is the weight of the synaptic connection from inhibitory neuron i to excitatory neuron j.

Excitatory weights are bounded so that their values lie between 0 and *w*_max_, and inhibitory weights are bounded so that they lie between −*w*_max-inh_ and 0. Note that the competitive nature of the BCM learning rule usually causes a single synapse to ‘win’, while all remaining synapses onto a neuron are pushed down to their minimum weight after a winning synapse emerges. This effect can be counteracted by setting a sufficiently low maximum weight, so that multiple synapses with large weights are required in order to achieve the target firing rate y_0_. The maximum synaptic weight *w*_max_ is set such that the target rate of a neuron will be achieved when a subset of synapses saturate at their maximum value, and another subset reach an intermediate synaptic weight. In this way multiple stable synaptic weights corresponding to multiple levels of co-activity can be achieved with the BCM learning rule.

The homeostatic target, y_0_, is the same for both inhibitory plasticity and the homeostatic component of BCM plasticity. Note that the speed of both learning rates *α* and η are artificially increased in order to reduce the computational resources required to simulate our network model. The timescales of synaptic plasticity in our network models are in the order of hundreds of seconds, while synaptic plasticity responses to visual deprivation occur over the course of days in vivo. This increased learning rate does not qualitatively affect our results, as there is a sufficient separation of timescales between synaptic plasticity and network dynamics.

### Seeding spontaneous subnetworks

We randomly seed spontaneous subnetworks within the initial synaptic weight matrix of the recurrent network model. This is achieved by randomly assigning each excitatory neuron to a spontaneous subnetwork, and then setting all weights between neurons in the same subnetwork to *w*_max_. The rest of the weights in the network are set to 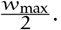 Note that this spontaneous subnetwork structure could also be achieved through Hebbian plasticity in combination with correlated spontaneous activity.

For uniform spontaneous subnetwork sizes, there are 20 neurons in each subnetwork. For diverse subnetwork sizes, the large subnetworks have 140 neurons and the remaining small subnetworks have 20 neurons.

### Simulating development and visual deprivation

Visual stimulus preference is assigned to excitatory neurons, ordered by neuron indices. This ensures that each neuron is independently a member of a visual and spontaneous subnetwork. There are 10 visual stimuli in the network, with 40 neurons per stimulus (neurons 1-40 prefer the first stimulus, neurons 41-80 prefer the second stimulus, etc.).

Every 500 ms, a randomly chosen visual stimulus is presented to the network. Neurons which prefer this visual stimulus receive a large external input with a firing rate of 12 Hz, while the remaining neurons in the network receive external input with a lower baseline firing rate of 4 Hz. These firing rates are kept constant for the next 500 ms. Throughout the entire simulation each neuron also receives independent external noisy inputs, modelled as an Ornstein-Uhlenbeck process with zero mean, standard deviation of 5 Hz, and time constant of 50 ms [Ricciardi and Sacerdote, 1979].

To simulate visual deprivation, we stop presenting visual stimuli to the network. External visual input to the neurons remains absent throughout deprivation, since monocular enucleation is a non-reversible visual deprivation paradigm. The inhibition-excitation (I/E) ratio was measured by calculating the ratio of incoming inhibitory synaptic currents to excitatory synaptic currents for each neuron.

### Measuring persistent correlations in the network model

We measure correlations by calculating the Pearson correlation coefficient of firing rate time series for each pair of excitatory neurons in the network. These measurements are performed on the last 500s of simulated activity after both development and visual deprivation. Two neurons are defined to be persistently correlated if their Pearson correlation coefficient is statistically significant and positive both after development and after the homeostatic response to visual deprivation. Neurons have a low proportion persistent correlations if they are persistently correlated with < 15% of other neurons, whereas neurons have a high proportion of persistent correlations if they are correlated with > 15% of other neurons. The normalised recovery given by 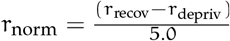, where r_recov_ is the firing rate after recovery, rdepriv is the firing rate immediately after visual deprivation, and 5.0 is the homeostatic target firing rate.

### Measuring persistent correlations in vivo

Data from experiments conducted by [Barnes et al., 2015] were analysed using a measure of persistent correlations. Briefly, we perform chronic imaging of calcium signals in layer 2/3 cells in monocular visual cortex of adult mice. Genetically encoded calcium indicator GCaMP_5_ signals were recorded with cellular resolution in behaving mice on a spherical treadmill [Akerboom et al., 2012, Keck et al., 2013]. Visual deprivation was performed via monocular enucleation, and cells were imaged for the 24 hours before and 72 hours after visual deprivation. Cells were classified into putative excitatory and putative inhibitory neurons using their calcium transients kinetics and immunohistochemistry (see [Barnes et al., 2015]). Only cells with significant responses to visual stimuli prior to visual deprivation and which exhibited some degree of recovery from visual deprivation were included in the analysis. The Pearson correlation coefficient of calcium traces were measured for each pair of putative excitatory neurons, while the mouse was stationary and in a dark room. This state is used as a proxy for spontaneous activity. Cells with a positive and statistically significant Pearson correlation coefficient are assumed to be correlated (p < 0.05). 252 correlated pairs of cells were observed prior to visual deprivation amongst 35 neurons, pooled across 4 animals. Persistent correlations are those which are present between cells both before visual deprivation, and 72 hours after visual deprivation. As with analysis of the network model, cells have a low proportion persistent correlations if they are persistently correlated with < 35% of other cells (N=15), whereas cells have a high proportion of persistent correlations if they are correlated with > 35% of other cells (N=20). This cut-off is higher than that used in the network model, as the average amount of persistent correlations is higher in vivo compared with the model. The normalised recovery is the ratio of the mean calcium trace at the end of the simulation compared with the mean calcium trace immediately prior to visual deprivation, for each cell. See [Keck et al., 2013, Barnes et al., 2015] for further details of experimental procedures.

### Simulating synaptic scaling

We simulate the effect of synaptic scaling using a simplification of multiplicative synaptic scaling. The dynamics of synaptic scaling are well characterised in previous theoretical studies; as such as we only simulate the overall weight changes after a prolonged period of synaptic scaling [Tetzlaff et al., 2011, van Rossum et al., 2000]. To this end we simply multiply all recurrent excitatory synaptic weights weights by a scaling factor β,

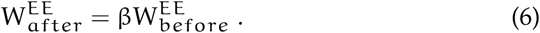

Visual deprivation is simulated as before, and weights are changed after visual deprivation. Weights are kept unbounded, so a synaptic weight that was *w*_max_ before scaling is β*w*_max_ after scaling. We chose β = 1.5, since it drives a homeostatic recovery to pre-deprivation firing rates. Decreasing β leads to a weaker recovery, and increasing β leads to firing rates above the pre-deprivation level.

### Model implementation and parameters

Parameters used to implement the simulations are shown in Table 1. Parameters for the neurons model are similar to those in previous studies [Rajan et al., 2010, Hennequin et al., 2014]. The choice of *w*_max_ = 0.015 is primarily determined by the constraint of exhibiting reasonable firing rates both before and in response to visual deprivation, and by the size of the network being simulated. *w*_max_ must be high enough such that neurons receive excitation primarily through recurrent inputs (as observed in cortical networks), but also low enough such that changes in feedforward inputs elicit dynamic network responses. The plasticity rates, *α*, η and τ_θ_ were chosen to be slow enough that there was a separation of timescales between firing rate dynamics and synaptic weight dynamics, thus avoiding oscillatory behaviour [Harnack et al., 2015].

**Table 1:** Simulation Parameters.

|  |  |  |  |  |  |  |  |
| --- | --- | --- | --- | --- | --- | --- | --- |
| $N_E$ | 400 | $N_I$ | 100 | $r_0$ | 1.0 Hz | $r_{\max}$ | 20.0 Hz |
| $dt$ | 0.05 ms | $\alpha$ | $2.0 \times 10^{-6}$ Hz | $\tau_\theta$ | $5 \times 10^4$ ms | $\eta$ | $5.0 \times 10^{-6}$ Hz |
| $w_{\max}$ | 0.015 | $w_{\max\text{-inh}}$ | 0.16 | $y_0$ | 5 Hz | | |

Numerical integration of Equation 2 through to Equation 5 was performed using the Euler method with a timestep of dt = 0.05 ms. Network rate dynamics were allowed to stabilise for 5000 ms with static synaptic weights before simulating development and visual deprivation. Network simulations and data analysis was performed with the numpy Python package and plotting with the matplotlib package and IPython notebook. [Hunter, 2007, van der Walt et al., 2011, Perez and Granger, 2007]. Code will be made publicly available on github and modeldb, and can be made available to reviewers [Hines et al., 2007].

## RESULTS

### Neurons in large spontaneous subnetworks recover from visual deprivation

Our network is composed of excitatory and inhibitory rate-based neurons with all-to-all recurrent synaptic connectivity (Figure 1A,B). The initial synaptic weight matrix is determined by seeded spontaneous subnetworks of different sizes (Figure 1C,D). Each excitatory neuron receives input corresponding to a given visual stimulus. As such, neurons are driven strongly both by visual inputs, and by neurons which are part of the same spontaneous subnetwork. Local connectivity is developed by presenting random sequences of visual stimuli to the network. Synaptic weights undergo plasticity as neurons which are assigned the same visual stimulus preference become coactive (Figure 1F). Synaptic weights below a defined threshold are removed (Figure 1F).

**Figure 1:**
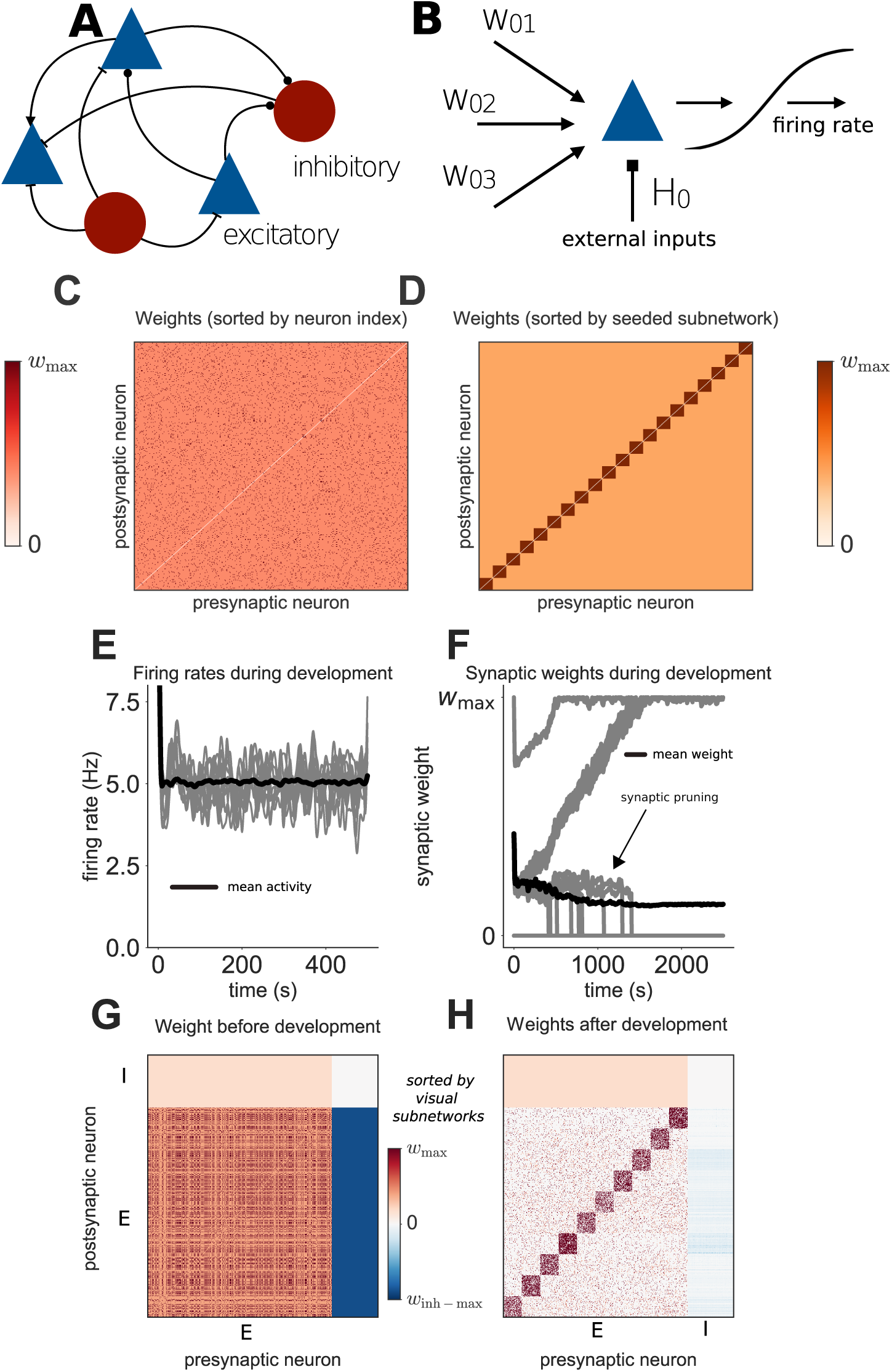
Simulating development in a network with uniform spontaneous subnetworks. **(A)** Architecture of the recurrent network model. **(B)** Neurons receive input from recurrent and external sources. **(C)** Initial excitatory synaptic weight matrix of an example network. **(D)** Same as **C**, sorted by grouping neurons into their seeded subnetworks. An orange colourmap is used throughout this paper to indicate sorting by seeded subnetworks, whereas a red colourmap is used to indicate sorting by visual subnetworks. **(E)** Evolution of individual (grey) and mean (black) firing rates of excitatory neurons during development. **(F)** Evolution of individual (grey) and mean (black) synaptic weights over time. Vertical grey lines indicate individual synapses being pruned. **(G)** Synaptic weight matrix before development. **(H)** Synaptic weight matrix after development. E and I refer to excitatory and inhibitory neurons, respectively.

Subnetworks of strongly interconnected neurons which share the same visual stimulus preference clearly emerge, due to neurons within networks receiving correlated external input (Figure 1H). This is in agreement with previous experimental and theoretical observations [Clopath et al., 2010, Ko et al., 2013, Ko et al., 2014]. The spontaneous subnetworks remain strongly interconnected (Figure 1G,H), with synaptic weights larger than the average synaptic weight in the network. Synapses are gradually pruned between neurons which do not form either a visual or spontaneous subnetwork together (Figure 1F,H).

We next simulate visual deprivation by turning off external inputs corresponding to visual stimuli, similar to previous models of visual deprivation [Intrator and Cooper, 1992, Toyoizumi et al., 2014]. The activity of neurons within different subnetworks clearly drops when visual input is removed (Figure 2A). Activity does not recover following deprivation, even though inhibitory drive is also reduced (Figure 2A). The final synaptic weight matrix illustrates this reduction in inhibition which occurs after visual deprivation (Figure 2B, compared with Figure 1H). These results suggest that there is not enough spontaneous recurrent excitation within small spontaneous subnetworks to drive a sufficient recovery from visual deprivation in the absence of synaptic scaling, despite reduced inhibition.

**Figure 2:**
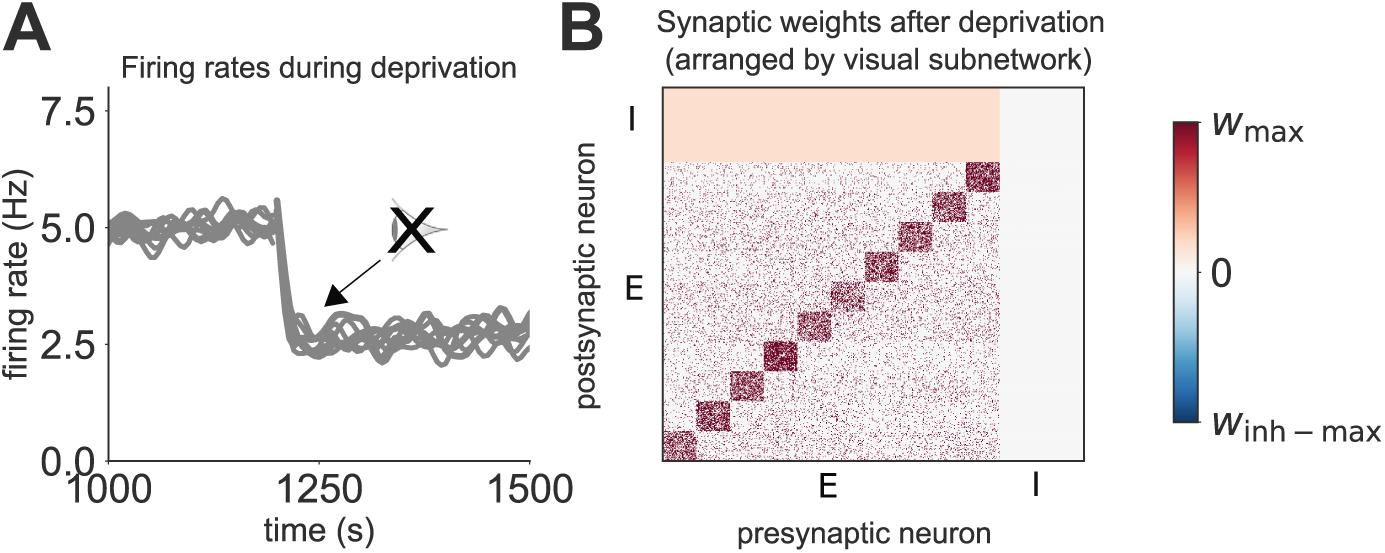
No recovery from visual deprivation in a network with uniform spontaneous subnetworks. **(A)** Network activity in response to deprivation, grouped into the different spontaneous subnetworks, each composed of 20 neurons (each grey line corresponds to a single subnetwork). **(B)** Synaptic weight matrix after visual deprivation. E and I refer to excitatory and inhibitory neurons, respectively.

We now test whether introducing diversity in the spontaneous subnetwork size influences the recovery of these subnetworks from visual deprivation. Since the stronger recurrent excitatory drive within larger subnetworks will recruit more inhibition to these subnetworks during the developmental phase of our simulation, the disinhibitory response to visual deprivation will have a larger range, and may be sufficient to unmask the spontaneous recurrent excitation within these larger spontaneous subnetworks. We therefore hypothesise that large spontaneous subnetworks will exhibit a larger homeostatic response to visual deprivation than small spontaneous subnetworks.

In order to test this hypothesis we first simulate the development of visual stimulus selectivity within networks with diverse sizes of seeded spontaneous subnetworks (Figure 3A). We designate two large subnetworks (coloured grey), while the remaining subnetworks are small (coloured black, see Methods). As visual stimuli to the network have not changed, the structure of the strongly interconnected subnetworks corresponding to these visual stimuli also remains unchanged (Figure 3B,E, compared with Figure 1H). There is however a difference in the amount of inhibition within the network after development and pruning. Since there is more recurrent excitation due to the large spontaneous subnetwork, there is also stronger inhibition before deprivation at the large spontaneous subnetwork (Figure 3E compared with Figure 1H).

**Figure 3:**
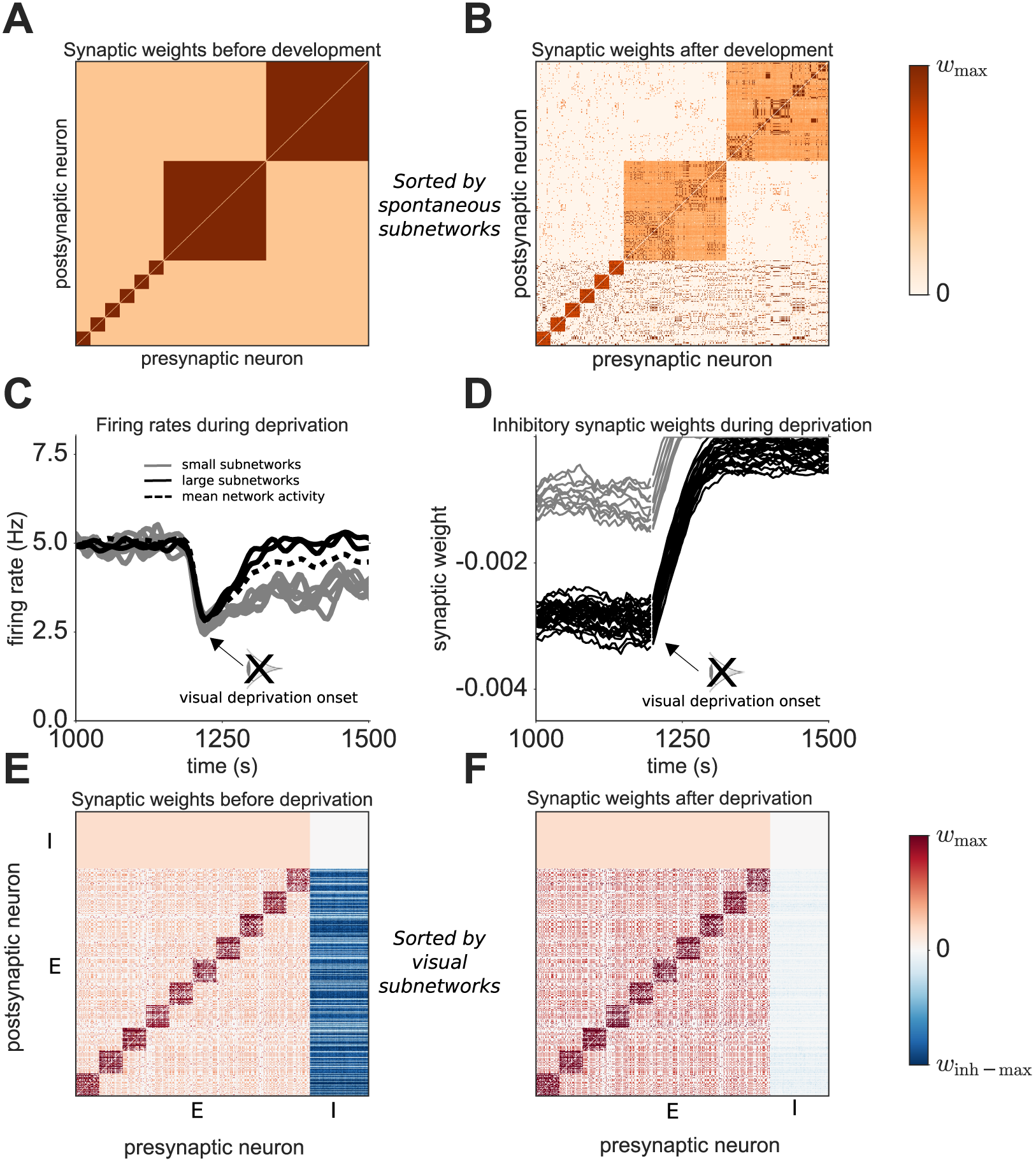
Recovery from visual deprivation in a network with diverse spontaneous subnetworks. **(A)** Initial excitatory weight matrix of a network with diverse seeded spontaneous subnetworks. Neurons are ordered by their spontaneous subnetworks membership. **(B)** Excitatory weight matrix of the network after development. Neurons are ordered by their spontaneous subnetworks membership. **(C)** Network activity in response to deprivation for small (grey) and large (black) spontaneous subnetworks. Dotted black line indicates mean network activity. **(D)** Changes in inhibitory synaptic weights in response to deprivation, for randomly chosen sample weights which connect to either small (grey) or large (black) spontaneous subnetworks. **(E)** Weight matrix before visual deprivation, sorted by visual subnetwork. **(F)** Weight matrix after visual deprivation, sorted by visual subnetwork. E and I refer to excitatory and inhibitory neurons, respectively.

As before (Figure 2), neurons in small spontaneous subnetworks do not recover from visual deprivation. In contrast, neurons embedded in large subnetworks exhibit a recovery from visual deprivation, returning to their pre-deprivation activity (Figure 3C, black lines). These results suggest that a subset of neurons will recover partially from visual deprivation and that a key factor in this recovery is the size of the network to which the neuron belongs. In agreement with experimental findings, the diverse homeostatic responses emerge despite blanket disinhibition occurring across the entire network [Barnes et al., 2015]. Since both recovering and non-recovering responses to visual deprivation were observed amongst neurons in different subnetworks in vivo, the dependence of recovery on a neuron’s subnetwork size provides a parsimonious explanation for this observation [Barnes et al., 2015]. As strong recurrent excitatory drive within large subnetworks recruits more inhibition to neurons in these subnetworks during development, the disinhibitory response of these neurons to visual deprivation will have a larger range. Disinhibition can therefore unmask sufficient spontaneous recurrent excitation within large spontaneous subnetworks to drive homeostatic recovery of activity, but not within small spontaneous subnetworks. As a consequence, neurons with identical membrane properties and firing rate statistics prior to deprivation can exhibit distinct homeostatic responses to visual deprivation by belonging to different spontaneous subnetworks.

### Visual deprivation reduces inhibition-excitation ratio across the network

[Barnes et al., 2015] crucially found a persistent reduction in the inhibition-excitation ratio following visual deprivation. We can test whether our model reproduces this observation by measuring the inhibition-excitation ratio before and after simulating visual deprivation (Figure 4A). As observed experimentally, there is a significant decrease in the inhibition-excitation (I/E) ratio after visual deprivation (grey bar) compared with the ratio immediately before visual deprivation (black bar, see Methods). The low I/E ratio is a consequence of the reduction in inhibitory synaptic weights which occur during the homeostatic recovery of a subset of neurons within the network.

**Figure 4:**
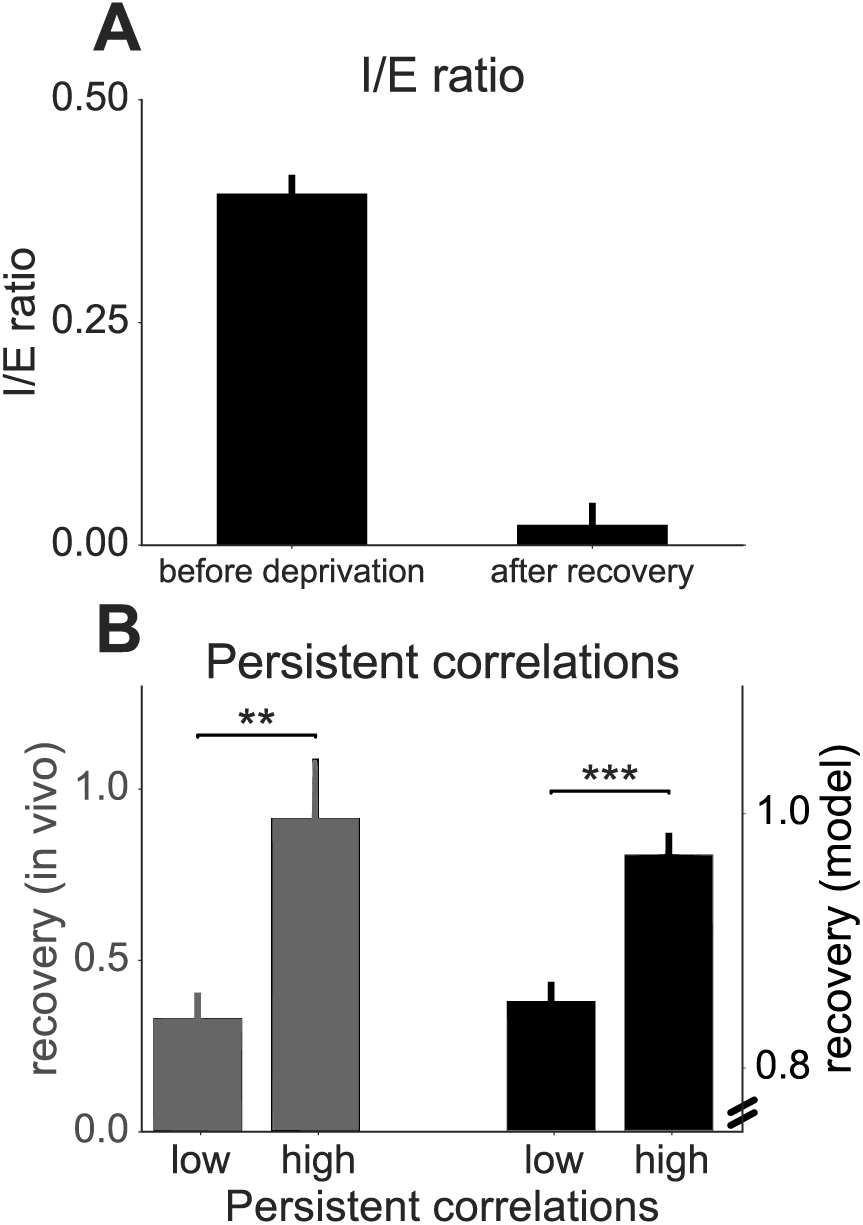
Experimental validation of the model. **(A) Reduction of inhibition-excitation ratio after visual deprivation** Ratio of incoming inhibitory to excitatory current for excitatory neurons in the network before visual deprivation and after partial recovery from visual deprivation. **(B) Model predictions are validated experimentally.** Persistence of correlations predicts the degree of recovery after visual deprivation in the network model and in vivo. Normalised recovery from visual deprivation for neurons with either a low or high proportion of persistent correlations, observed in vivo (grey bars) and in our network model (black bars). Note the different y-axis scale when comparing in vivo and network model results. ** p < 0.01, *** p < 0.001, independent 2-sample t-test. Error bars in both plots indicate the standard error of the mean.

### The size of enduring functional networks predicts homeostatic recovery

Our network model predicts that neurons belonging to large functional networks that endure throughout deprivation will exhibit the strongest homeostatic recovery (Figure 4B, see Methods). In order to test this prediction empirically, we used data from calcium imaging experiments of L2/3 excitatory neurons in adult mouse V1 (see methods). L2/3 excitatory neurons were chronically measured both before and after (+72 hours) monocular enucleation.

To estimate the size of functional networks in vivo we measured correlations between calcium signals from the imaged L2/3 neurons. Neuronal calcium correlations have been shown to represent either reciprocal connections or common inputs in numerous cortical imaging experiments [Miller et al., 2014, Ko et al., 2014, Barnes et al., 2015].

At each neuron we compared the size of persistent functional networks which endured throughout deprivation to the degree of homeostatic recovery measured at 72 hours after enucleation. As predicted by our model, neurons with a high proportion of persistent correlations exhibit a strong homeostatic recovery from visual deprivation, suggesting that this recovery is enabled by spontaneous subnetworks which endure following visual deprivation (Figure 4B, see Methods). Neurons with a low proportion of persistent correlations exhibit very weak recovery. We propose that neurons with high persistent correlations receive a high proportion of recurrent excitation which is spontaneously driven as opposed to visually evoked, therefore driving a stronger homeostatic recovery from visual deprivation.

In summary, the predictions of our model are validated by experimental data, suggesting that the size of a functional network which a neuron is embedded in before deprivation can influence the strength of its homeostatic recovery.

### Synaptic scaling is not consistent with subnetwork-specific recovery

While evidence of excitatory synaptic scaling in response to visual deprivation has not been observed in layer 2/3 of adult mouse visual cortex [Barnes et al., 2015], it is a ubiquitous response to sensory deprivation in young animals [Desai et al., 2002], and has been observed in layer 5 of adult mouse visual cortex [Keck et al., 2013]. We therefore tested whether subnetwork-specific homeostatic responses can be explained by synaptic scaling alone. We do this by assuming a simplified form of synaptic scaling, in which all excitatory weights are multiplicatively increased in response to visual deprivation and inhibitory weights are kept fixed (Materials and Methods). Synaptic scaling leads to a full homeostatic recovery (Figure 5A). Crucially, this recovery occurs uniformly across all neurons, and as a consequence is not subnetwork-dependent. The I/E ratio of the network is only slightly reduced after recovery by synaptic scaling (Figure 5B), in contrast with the large reduction observed in our subnetwork-specific network model and *in vivo* (Figure 4A). Since all neurons recover uniformly in the network with synaptic scaling, we find no substantial difference in recovery strength between neurons with low or high persistent correlations (Figure 4C), again in contrast with observations from our subnetwork-specific model and *in vivo*.

**Figure 5:**
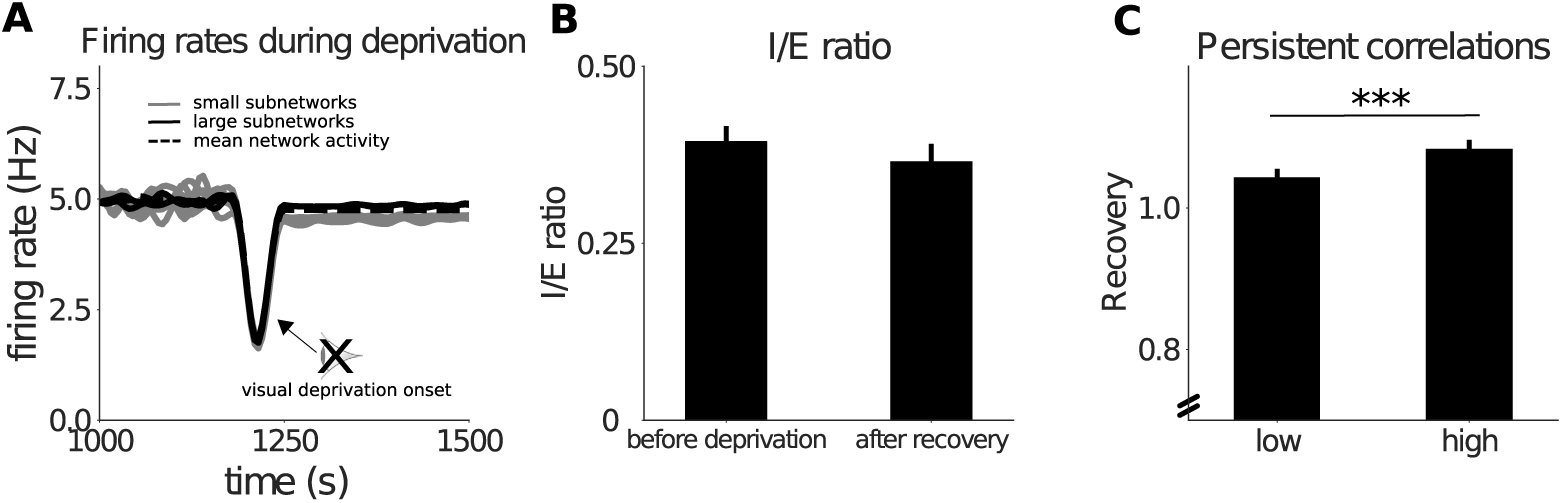
Synaptic scaling is not consistent with experimental observations. **(A) Uniform homeostatic responses mediated by synaptic scaling in a network with diverse spontaneous subnetworks.** Network activity in response to deprivation for small (grey) and large (black) spontaneous subnetworks. Dotted black line indicates mean network activity. **(B) Inhibition-excitation ratio is not substantially reduced after visual deprivation.** Ratio of incoming inhibitory to excitatory current for excitatory neurons in the network before visual deprivation and after recovery from visual deprivation by synaptic scaling. **(C) Persistence of correlations do not substantially determine homeostatic responses.** Normalised recovery from visual deprivation by synaptic scaling for neurons with either a low or high proportion of persistent correlations. *** p < 0.001, independent 2-sample t-test. Error bars in both plots indicate the standard error of the mean.

## DISCUSSION

Using a recurrent network model, we show that a combination of biologically plausible synaptic plasticity mechanisms can capture both the experience-dependent development of visual and non-visual spontaneous subnetworks in V1 as well as diverse subnetwork-specific homeostatic responses to visual deprivation. We find that the strength of homeostatic recovery for each neuron is subnetwork-specific. Neurons in large spontaneous subnetworks recover partially from visual deprivation, whereas neurons in small spontaneous subnetworks do not recover. This provides a possible explanation for the observed emergence of diverse activity profiles following visual deprivation in adult mice, despite blanket disinhibition occurring across all neurons [Barnes et al., 2015].

The mechanisms in our network model consist of Hebbian excitatory synaptic plasticity with activity-dependent control of the threshold for LTP over LTD (a BCM learning rule), and a homeostatic form of inhibitory plasticity which aims to maintain a target firing rate. We simulated development within the network such that strongly interconnected subnetworks are formed either by common visual input, or by correlated spontaneous activity within non-visual ‘seeded’ subnetworks (Figure 1). We could then study the effects of visual deprivation on either of these subnetwork types.

Visual deprivation leads to a reduction in activity, followed by homeostatic recovery amongst a subset of neurons (Figure 2, Figure 3). As hypothesised, this recovery is mediated by disinhibition, which unmasks the non-visual, spontaneously-driven subnetworks. Despite a subset of neurons exhibiting no homeostatic response, all neurons within the network underwent blanket disinhibition, reproducing the phenomenon observed by [Barnes et al., 2015]. Crucially, the strength of homeostatic recovery for each neuron is subnetwork-specific, and is determined by the size of the spontaneous subnetworks: neurons within small subnetworks exhibit very little recovery after deprivation (Figure 2), whereas neurons within large subnetworks recover almost completely (Figure 3). The inhibitory synaptic plasticity rule ensures that neurons within large spontaneous subnetworks recruit stronger inhibition during development in order to maintain firing rates similar to neurons within small subnetworks (Figure 3). As such, the homeostatic recovery observed amongst large subnetworks is not simply a consequence of neurons within these subnetworks receiving more excitatory connections. If this were the case, size-dependent differences in the spontaneous subnetwork responses to visual deprivation would be evident immediately after deprivation, rather than after a period of disinhibition. Our results instead demonstrate that neurons within large spontaneous subnetworks have increased potential for excitatory recurrent activity which can be unmasked through disinhibition.

We would not expect such a difference in the homeostatic response if we increased the number of synapses within visual subnetworks, as they will no longer be activated following visual deprivation. Similarly, spontaneous subnetworks do not share common visual inputs, so recovering neurons have independent visual stimulus preferences prior to visual deprivation - as observed by [Barnes et al., 2015]. Although there in no synaptic scaling in our network model, we show separately that a simple model of multiplicative synaptic scaling is sufficient to drive a full homeostatic recovery (Figure 5A). However, this recovery is not subnetwork-dependent, and occurs uniformly across all neurons. In our model however, we observed a reduction in the I/E ratio following visual deprivation (Figure 4A), consistent with experimental observations from [Barnes et al., 2015]. Although the I/E ratio reduced slightly when we simulated synaptic scaling, this was a substantially smaller reduction compared with that observed in our subnetwork-specific model (Figure 5B). Taken together, our model demonstrates that synaptic scaling is not required in order to trigger a homeostatic response to visual deprivation, and that inhibitory plasticity combined with adequate recurrent excitation is sufficient to drive a homeostatic response. This is in agreement with experimental observations, and provides an alternative to previous theoretical models of homeostatic recovery which propose synaptic scaling as the primary homeostatic mechanism [Barnes et al., 2015, van Rossum et al., 2000, Toyoizumi et al., 2014].

The dependence of the degree of homeostatic recovery on the size of spontaneous subnetworks provides a parsimonious explanation for the diverse and subnetwork-specific homeostatic responses described by [Barnes et al., 2015]. However, it remains unclear whether this form of homeostatic response is functionally advantageous compared with excitatory synaptic scaling, which drives a complete homeostatic recovery across all neurons (Figure 5A). An intriguing possibility is that, while synaptic scaling is arguably more effective at maintaining overall network activity, disinhibitory unmasking of recurrent excitation allows a network to conserve its underlying functional structure. This is supported by the observation that subnetwork-specific responses can reflect previously seeded subnetworks, and by the persistence of correlations observed both *in vivo* and in our subnetwork-specific network model. The conservation of subnetwork structure may be particularly important amongst L2/3 neurons, whose recurrent connections can develop strong functional ensembles [Ko et al., 2013].

Finally, our model predicts that neurons with high persistent correlations were more likely to exhibit a strong homeostatic recovery. This is because correlations within visual subnetworks are reduced by visual deprivation, whereas correlations within spontaneous subnetworks persist after visual deprivation, since they are unmasked by disinhibition. As such, neurons which receive more recurrent drive from spontaneous subnetworks exhibit both a stronger homeostatic recovery, and more persistent correlations (Figure 4B, black bars). Simulating homeostatic recovery through synaptic scaling alone leads to a markedly reduced dependence of recovery strength on persistent correlations compared with our subnetwork-specific model - largely because homeostatic recovery is uniform in the case of synaptic scaling (Figure 5C). We tested the prediction of our subnetwork-specific model experimentally, by measuring persistent correlations in calcium signals across cells in monocular visual cortex of adult mice throughout monocular enucleation. While calcium imaging cannot directly probe connectivity, cell-to-cell calcium signal correlations provide us with a reliable indicator of the degree of functional association between cells [Miller et al., 2014, Ko et al., 2014, Barnes et al., 2015]. We found that cells with a higher proportion of persistent correlations exhibited a stronger recovery from visual deprivation, verifying our prediction (Figure 4B, grey bars).

Our measure of persistent correlations cannot untangle the contributions to correlations mediated by common inputs from the contributions mediated by recurrent connectivity [Pernice et al., 2011, Doiron et al., 2016]. However, there is growing evidence that recurrently connected neurons in visual cortex are likely to share common inputs, indicating an interdependence in these two sources of correlations [Komiyama et al., 2010, Ko et al., 2013, Miller et al., 2014, Ko et al., 2014]. The importance of persistent correlations in predicting whether neurons recover suggests that homeostatic recovery is determined by pre-existing excitatory connections which endure and are functional following visual deprivation, and does not depend on the absolute number of excitatory recruitment or reorganisation of excitatory connections.

Although our network model suggests that subnetwork-specific recovery is size-dependent, the data do not rule out alternative theories of spontaneously-driven recovery. For example, diversity in the strength of recurrent synapses within subnetworks may have a similar effect as the diversity in subnetwork size implemented within our network model. This could be due to stronger correlations within some subnetworks, which mediate increased synaptic potentiation during development. This would lead to diverse, subnetwork-specific homeostatic response which are independent of subnetwork size, but dependent on recurrent synaptic strength within subnetworks. A second possibility is that the degree of recovery for each neuron is determined not by specific subnetwork properties, but by the strength of spontaneous excitatory input currents which individual neurons receive prior to deprivation. As there is balanced excitatory and inhibitory drive in healthy cortical networks, increased recurrent input would not necessarily alter neural firing rates, but would ensure a larger dynamic range over which disinhibition mediates homeostatic recovery [Moreau et al., 2010, Dorrn et al., 2010]. This diversity of spontaneous drive in neurons has recently been experimentally observed in visual cortex [Okun et al., 2015].

There are a number of plausible mechanisms which may lead to the formation of seeded subnetworks prior to visual experience. Functional sub-networks can be formed through enhanced electrical coupling, observed between neurons derived from the same progenitor cell [Yu et al., 2012, Li et al., 2012]. Such a scenario has previously been explored in a computational network model, in which the development of diversely sized subnetworks was observed [Bauer et al., 2014]. Additionally, common input from non-visual modalities may lead to the formation of functional subnetworks, much in the same way that common visual inputs drives the formation of visual subnetworks [Ko et al., 2013, Miller et al., 2014]. These subnetworks may not be seeded as we described, but could form under the influence of Hebbian plasticity [Litwin-Kumar and Doiron, 2014]. Diverse homeostatic recovery could occur as observed with spontaneous subnetworks, provided that monocular visual deprivation preserved the correlated structure of inputs from these non-visual modalities. The multimodal response properties of V1 neurons observed in recent experiments suggest many potential non-visual sources, including sound-evoked inputs from auditory cortex, movement-related inputs, or top-down projections from higher cortical regions [Zhang et al., 2014, Ibrahim et al., 2016, Makino and Komiyama, 2015, Keller et al., 2012]. The subnetwork size is largely unconstrained by data. Subnetworks composed of as little as 10 neurons can form, as demonstrated through induced LTP by repeated triggering of spike trains in a randomly chosen group of L2/3 excitatory neurons [Kim et al., 2016]. Large-scale recording of neurons reveal ensembles in which 5 − 10% of imaged neurons are active during an ensemble event [Miller et al., 2014]. However, since these ensembles are extracted by measuring coactivity during a given time window, the sparse nature of cortical activity likely means that the actual number of neurons which form these ensembles is underestimated.

The network model we propose here remains agnostic to the origin of seeded, non-visual subnetworks. Nonetheless, our model demonstrates that the simultaneous development and maintenance of multiple subnetworks with different modalities is possible in a recurrent network using a simple combination of synaptic plasticity mechanisms. Moreover, neurons may flexibly participate in multiple subnetworks, as observed in primary visual cortex [Miller et al., 2014]. Disinhibitory unmasking of these flexible subnetworks after visual deprivation, as demonstrated in our model, may form the basis of extensive crossmodal plasticity observed in adult visual cortex [Lee and Whitt, 2015, Nys et al., 2014, VanBrussel et al., 2011]. Further experiments may uncover the multimodal nature of these diverse subnetworks.

## AUTHOR CONTRIBUTIONS

Y.S., S.J.B. and C.C. conceived and designed experiments, and wrote the manuscript. Y.S. developed computational network model. S.J.B. collected and analysed imaging data.

## ACKNOWLEDGEMENTS

We thank Tara Keck and Georg Keller for generously sharing their data, and for their invaluable comments.

